# Methods for Quantitative Analyses of Nerve Fiber Deformation in the Myenteric Plexus Under Loading of Mouse Distal Colon and Rectum

**DOI:** 10.1101/2025.04.02.646858

**Authors:** Amirhossein Shokrani, Atta Seck, Bin Feng, David M. Pierce

## Abstract

Visceral pain in the large bowel is a hallmark of irritable bowel syndrome (IBS) and the primary reason patients seek gastroenterological care. Notably, mechanical distension of the distal colon and rectum (colorectum) reliably evokes abdominal pain and thus understanding mechanotransduction of sensory nerve endings (nerve fibers) in the colorectum is crucial for understanding and treating IBS-related bowel pain. To facilitate such understanding we aimed to establish novel methods to mechanically test, image, and analyze large-strain deformations of networks of nerve fibers in the myenteric plexus of the colorectum, and thus enable quantitative analyses. We successfully delivered circumferential deformation (force or displacement driven) to intact segments of colorectum while maintaining the myenteric plexus in focus during fluorescent imaging to capture the deforming nerve fibers. We also established a semi-automated method to recapitulate the network morphology and a code to calculate the stretch ratios of individual nerve fibers deforming within the myenteric plexus of mouse colorectum. Our code allows plotting of stretch ratios for each fiber, stretch ratios vs. fiber angle, and stretch ratios vs. fiber length. Our methods not only facilitate analyses of deformations of networks of colorectal nerve fibers in the context of visceral nociception but are also applicable to analyzing the in-plane deformation of other two-dimensional fiber networks. We provide free, public access to our analysis code for MATLAB, including input files for a simple test case, at github.uconn.edu/imLab/Fiber-Network_Analyses.

## 1. Introduction

Individuals with irritable bowel syndrome (IBS) frequently experience considerable discomfort due to visceral pain, which significantly diminishes their quality of life [1, 2]. This pain often originates in the lower abdominal area, particularly the distal colon and rectum (colorectum), presenting as a deep, dull, cramping sensation, in contrast to the sharp pain typical of cutaneous issues [2]. Addressing visceral pain in IBS poses a substantial clinical challenge. Many drugs effective against other pain types either fail to relieve visceral pain or cause severe gastrointestinal (GI) side effects [3].

Mechanotransduction of afferent nerve endings within the colorectum is crucial for visceral nociceptive perception [4, 5, 6]. In particular, stimuli such as cutting, pinching, and heating that trigger significant pain in skin do not produce pain responses in the GI tract. However, distension of the distal colon and rectum consistently induces visceral pain in healthy individuals and intensifies discomfort in IBS patients [4, 7, 8]. Enhanced understanding of colorectal mechanotransduction during deformation of the bulk tissue could significantly advance therapeutic strategies for IBS-associated visceral pain [6, 9].

The sensation of pain arises from activation of a population of sensory (afferent) neurons known as nociceptors, specifically attuned to encode injurious stimuli. Most stretch-sensitive colorectal afferents are considered nociceptors, comprising unmyelinated C-fibers responding to noxious levels of distension and able to undergo sensitization [1, 6]. Their role in detecting and transmitting pain signals underscores their importance in the pathophysiology of conditions like IBS, where mechanical distension of the colorectum is a key factor in the perception of pain. Understanding the mechanisms by which these nociceptors function and become sensitized is crucial for developing targeted therapies for visceral pain [2, 10, 11].

Recent anatomical studies utilizing optical tissue clearing carefully quantified afferent fiber density and morphology across various colorectal layers and revealed concentrated afferent fibers in the myenteric plexus and the submucosa [12, 13, 14, 15]. Results demonstrate that approximately 22% of colorectal afferents terminate within the ganglia of the myenteric plexus, an integral component of the intrinsic nervous system in the GI tract. Furthermore, afferent endings within the myenteric plexus potentially play crucial roles in visceral nociception [16, 17].

In the current study we aim to establish novel methods to quantify the biomechanics of deforming nerve fibers in the myenteric plexus under mechanical loading of colorectums of mice. Specifically, our methods will further advance mechanistic understanding of afferent mechanotransduction by quantitatively determining the stretch ratios of microns-thick nerve endings in the myenteric plexus during macroscopic colorectal distension. Such studies will help elucidate patterns of deformation of the nerve fibers and specifically facilitate comparison of these with heterogeneous mechanical strains within the myenteric.

## 2. Materials and Methods

### 2.1. Preparing the colorectal specimens

The University of Connecticut Institutional Animal Care and Use Committee (UConn IACUC) reviewed and approved all experiments on mice. Animal care services staff housed mice in pathogen-free facilities assured by the Public Health Service and accredited by the American Association for Accreditation of Laboratory Animal Care (AAALAC) following the Guide for the Care and Use of Laboratory Animals Eighth Edition. UConn is an AAALAC accredited institution, accreditation requiring extensive internal review and validation by AAALAC evaluators. Mice resided in individual ventilated caging systems in polycarbonate cages (Animal Care System M.I.C.E.) with a maximum of 5 animals per cage and provided with contact bedding (Envigo T7990 B.G. Irradiated Teklad Sani-Chips). Staff fed mice ad lib using either 2918 Irradiated Teklad Global 18% Rodent Diet or 7904 Irradiated S2335 Mouse Breeder Diet supplied by Envigo and supplied reverse-osmosis water chlorinated to 2 ppm using a water bottle. Nestlets and huts provided enrichment. Staff maintained rodent housing at 73.5^°^F (70 to 77^°^F) with humidity at 50% (35 to 65%) and on a 12:12 light-dark cycle. Animal care services staff observed animals daily and changed every two weeks.

To demonstrate our methods we utilized transgenic mice, heterozygous for VGLUT2-Cre and tdTomato (VGLUT2/tdT) genes of both sexes, aged between 18-35 weeks. The VGLUT2 promotor drives expression of tdTomato in most colorectal sensory afferent neurons [18, 15, 19]. We anesthetized mice by isoflurane inhalation (2–5%) until it lost plantar reflex to forceps pinching. Then, we performed euthanasia by transcardial perfusion with oxygenated Krebs solution (in mM: 117.9 NaCl, 4.7 KCl, 25 NaHCO_3_, 1.3 NaH_2_PO_4_, 1.2 MgSO_4_, 2.5 CaCl_2_, and 11.1 D-glucose) at room temperature and bubbled with 95% O_2_ and 5% CO_2_ from the left ventricle to the right atrium through the circulatory system. Subsequently, we harvested the distal 22mm of the colorectum with the mouse submerged in fresh oxygenated Krebs solution. Finally, we carefully removed connective tissues and the mesentery to create a flat specimen.

We use animals of both sexes equally and so sex did not influence the results from our methods.

### 2.2. Mechanical testing in uniaxial extension

We applied uniaxial, circumferential mechanical stretch to the intact, tubular colorectum via two intraluminal cylindrical rods of diameter 1.2 mm. The smooth stainless steel rods minimally impact the axial load (and deformation) of the colorectum, enabling focused delivery of circumferential stretch while maintaining a stable focal plane within the myenteric plexus (see fig. 1). Within a 3-D printed chamber, we positioned the colorectum, with the inserted rods, on a glass layer within the chamber to minimize friction. We then filled the chamber with oxygenated Krebs solution at room temperature to maintain the viability of the colorectum and embedded nerve fibers. We fixed one rod and actuated the other rod by pulling with a servo-controlled force actuator (Model 305D, Aurora Scientifics, Inc). To focus on the stress-strain response at relatively high intensity we pre-stretched the colorectum to 188.99 *±* 38.30mN and delivered a quasi-static, displacement-controlled ramp from 0 to 2 mm in 15 sec. We also pre-conditioned the specimen by repeating this ramped stretch 3-5 times before recording the force-displacement response to ensure repeatability [20].

**Figure 1.**
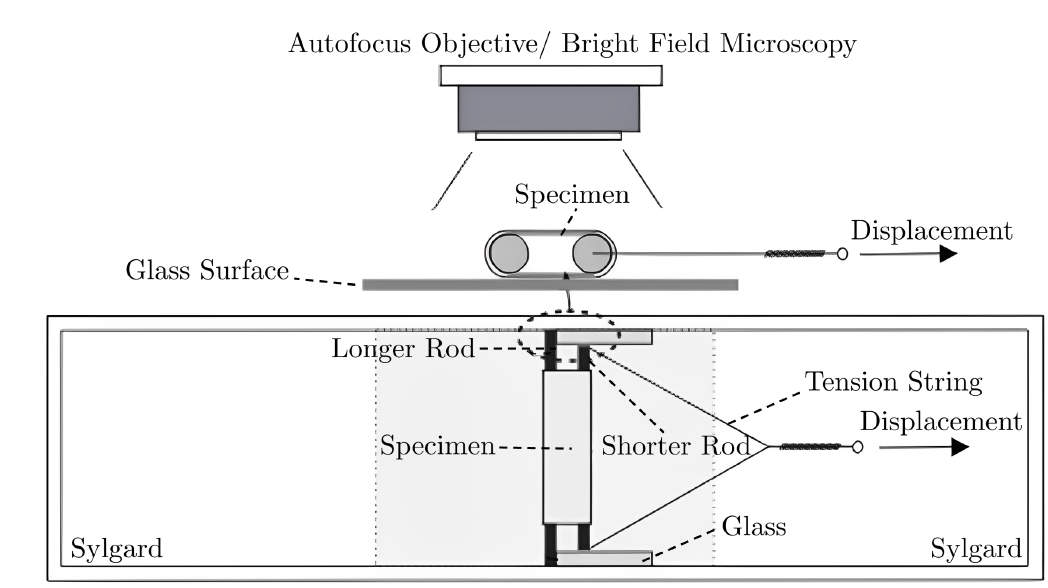
Schematic of the experimental setup for imaging deformation of a network of nerve fibers during circumferential, colorectal stretch. In a custom-built, 3-D-printed chamber we distended the colorectum using two intraluminal stainless-steel rods of lengths 25 mm and 30 mm. We fixed the longer rod while we applied stretch via the shorter rod. To minimize friction the entire assembly glides on the glass surfaces at the bottom and two sides of the chamber.

### 2.3. Fluorescent imaging during deformation

We visualized fluorescent nerve fibers within the VGLUT2/tdT mouse colorectum using an upright fluorescent microscope (BX51WI, Olympus, Inc) with a 4*×* objective lens (Model UPlan-FLN ∞/ - / FN26.5, NA 0.13, Olympus, Inc). Using the open-source microscopy software Micro-Manager [21], we imaged using a low-noise sCMOS camera (Zyla-4.2P, 82% quantum efficiency, Andor Technology) with high resolution (2048*×*2048 pixels) at ten frames per second. We captured the motion of VGLUT2-labeled unmyelinated nerve fibers during circumferential colorectal stretching using a field of view of 5.35*×*5.35 mm (383 pixels per mm). To image the colonic and rectal regions for both mesenteric and antimesenteric sides separately requires 2–4 cycles of quasi-static, ramped stretching. Throughout the procedure we observed no significant changes in the mechanical integrity of the colorectum.

### 2.4. Analyzing the images

We established a semi-automated framework to quantify the micron-scale stretch patterns with the network of nerve fibers within the myenteric plexus undergoing macroscopic circumferential stretch. Firsts we extract the fiber network from initial (reference) images of the colorectum. Then, our algorithm automatically tracks the deformation of the fiber network using images recorded from subsequent time points during the deformation and determines the stretch ratio of individual fiber bundles comprising the network. Finally, the algorithm generates plots of the stretch ratios as spatial heat maps and facilitates some quantitative analyses.

#### 2.4.1. Determining a point cloud

We utilized fluorescent images from VGLUT2-labeled unmyelinated nerve fibers within the myenteric plexus of mouse colorectum during circumferential stretch to capture the morphology and deformation of genetically labeled nerve fiber networks. First, we manually determined the coordinates of intersections among fiber bundles using ImageJ [22]. The intersection points are mostly the ganglia of the myenteric plexus in the intrinsic nervous system. Then, we identified points along in the fiber bundles (between the ganglia) where the deflection angle from straight was greater than 10.2 degrees. We determined such point clouds for three time steps of the experiment. We then wrote the complete point cloud as two space-delineated matrices of data: the first including point number (the row) and the *x* and *y* in-plane coordinates, and the second including the fiber connectivity, all for use in §2.4.2.

#### 2.4.2. Analyzing the network of nerve fibers

We established an algorithm in MATLAB (V2024, Mathworks, MA) to leverage the point-cloud information to process the data, calculate stretch ratios, and other relevant parameters, and overlay heat map data on the original images. To start the analyses we read the point coordinates at all time steps and calculating the Euclidean fiber lengths at each time step [23]. To quantify the deformation, we computed the stretch ratio for each fiber (defined by a pair of points) by normalizing the current fiber lengths by those determined in the initial or in the previous time step. In addition, we computed the angles between each pair of fibers originating from points where more that two fibers intersected, and we calculated the number of fibers connecting at each point. With this information we distinguished ganglia (connections among more than two fibers) from deflection points (connections among two segments of the same fiber). We then sorted and reported all calculated angles, both for all points (ganglia and deflection points) and specifically for points connecting exactly two fibers (deflection points). We utilized MATLAB’s image processing capabilities to overlay a heat map of the stretches within the fiber network onto the microscopy image of the network of fibers. Finally, we created plots of stretch ratios for each fiber, stretch ratios vs. fiber angle, and stretch ratios vs. fiber length.

## 3. Results

### 3.1. Imaging the colorectal specimen

In Fig. 2 we show a representative image of fluorescent nerve fibers from a VGLUT2/tdT mouse colorectum in the initial and final configurations.

**Figure 2.**
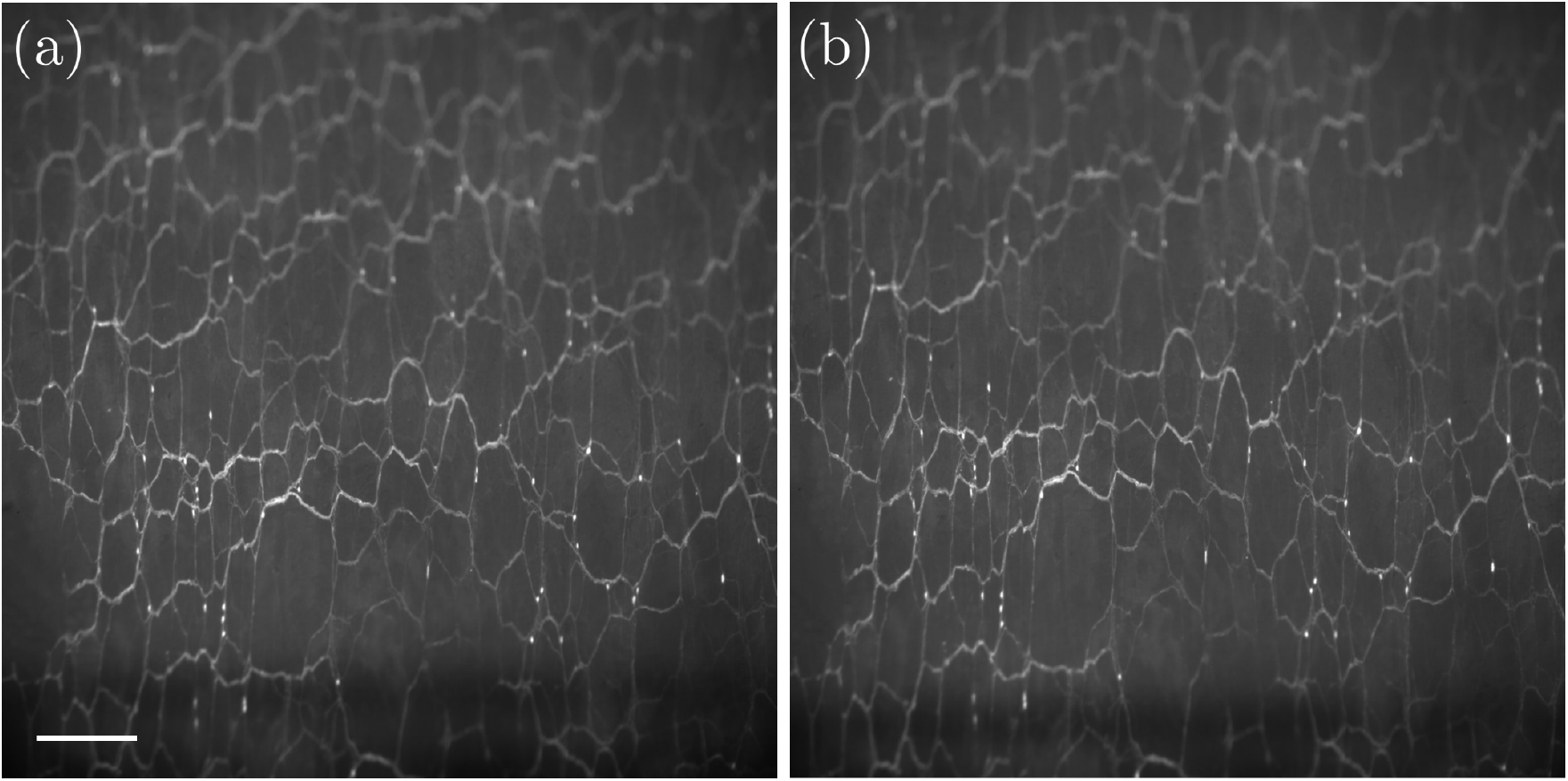
Representative magnified view of fluorescent microscopy images of VGLUT2-labeled nerve fibers during circumferential distension of the colon. (a) Image of initial configuration. (b) Corresponding image at maximum displacement. The scale bar is equal to 0.465 mm.

### 3.2. Analyzing the network of nerve fibers

In Fig. 3 we show a representative magnified image of fluorescent nerve fibers from a VG-LUT2/tdT mouse colorectum and the corresponding, overlayed map of fiber stretch ratios. Figure 3(a) shows a representative magnified view of fluorescent unmyelinated nerve fibers from the VGLUT2/tdT mouse colorectum with a field of view 3.57*×*3.57 mm, and a box highlighting the region for subsequent analyses. Figure 3(b) shows our heat map of fiber stretch ratios during circumferential colorectal stretching overlayed on the same microscopy image.

**Figure 3.**
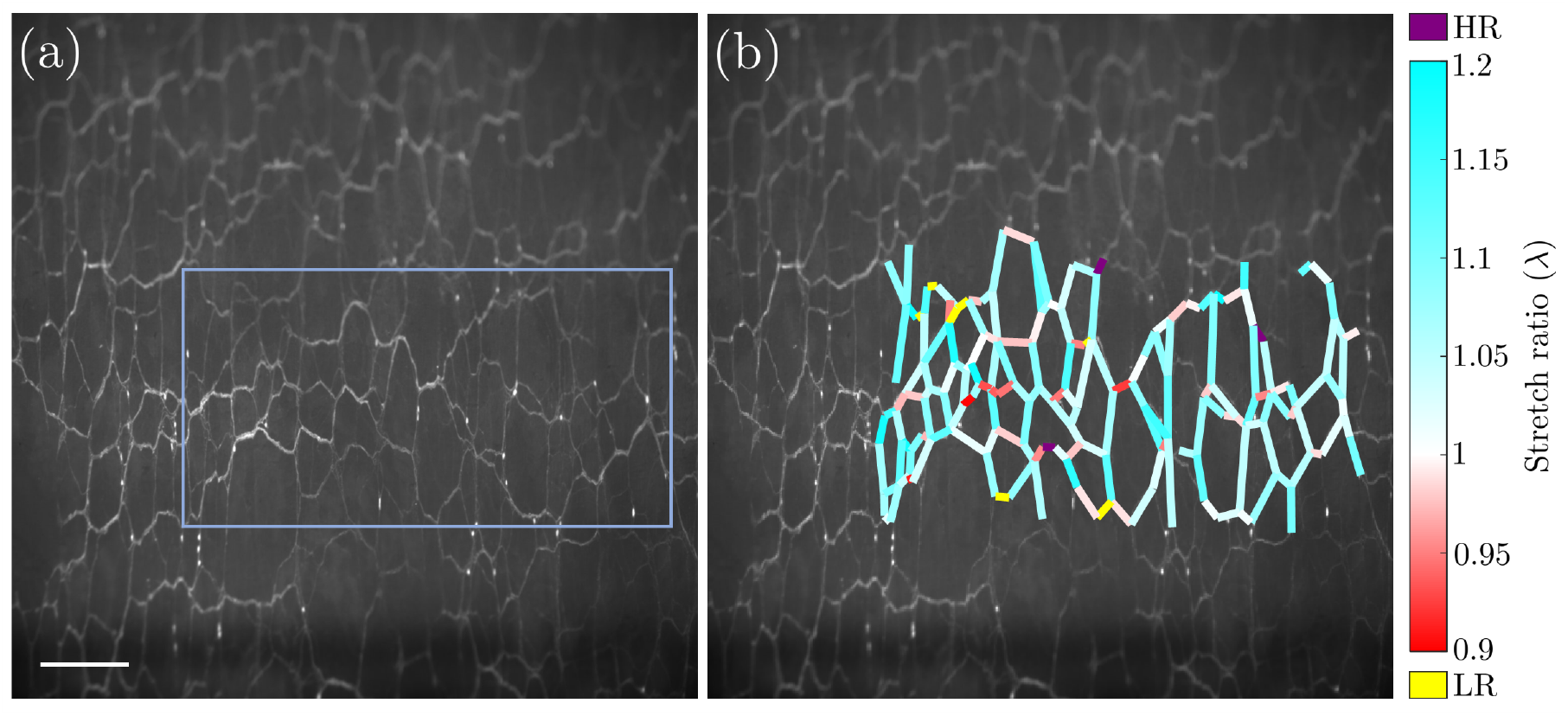
Representative magnified view of fluorescent microscopy images with corresponding analysis. (a) VGLUT2-labeled nerve fibers during circumferential distension of the colon with box highlighting area for subsequent analysis. (b) Overlay of fiber stretch ratios in highlighted area. HR: Higher range, stretch ratios above 1.2; LR: Lower range, stretch ratios below 0.9. The scale bar is equal to 0.465 mm.

In Fig. 4 we show the stretch ratio for each fiber and these stretch ratios with respect to the fiber’s angle with the *x*-axis (axial orientation within the colorectum).

**Figure 4.**
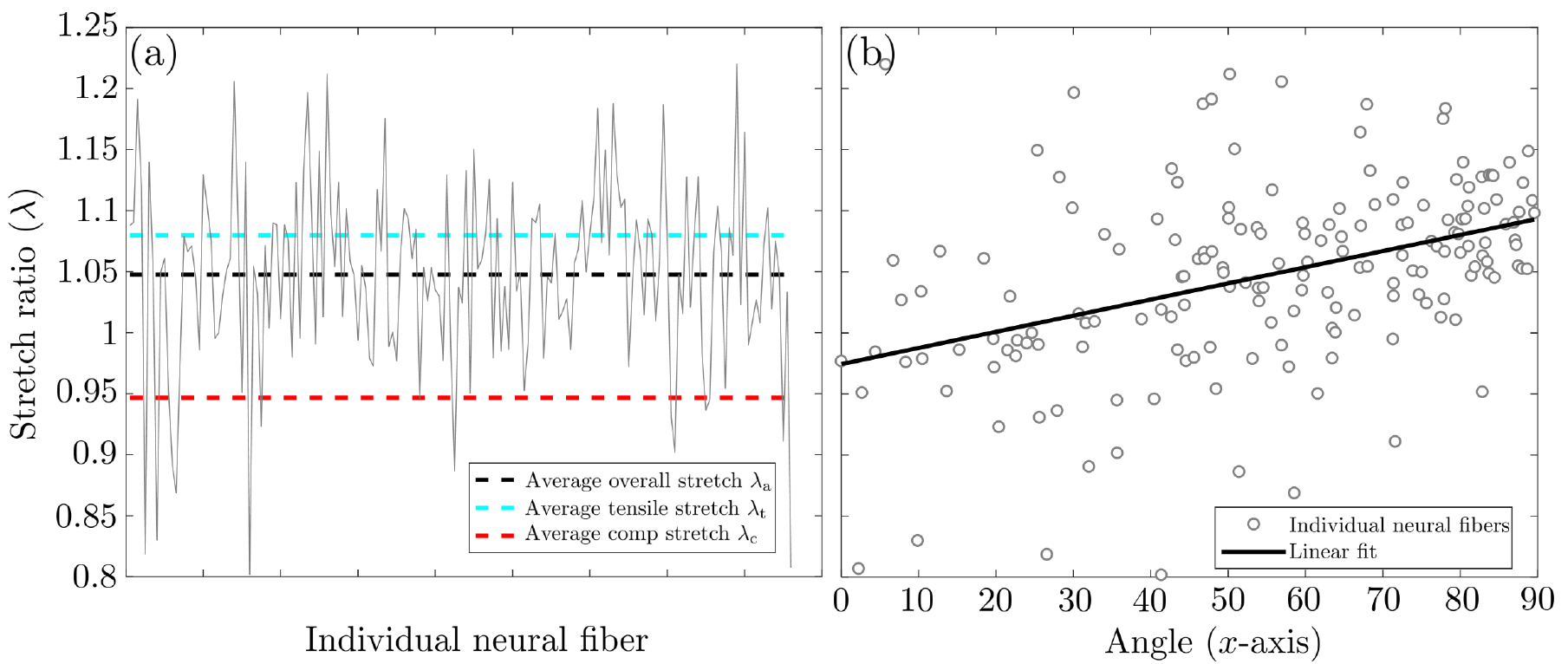
Representative stretch ratios of individual fibers during circumferential colorectal stretch. (a) Stretch ratios for individual fibers within the network analyzed. Black dashed line indicates average overall stretch ratio (*λ*_a_ = 1.048). Cyan and red dashed lines indicate the average tensile (*λ*_t_ = 1.080) and compressive (*λ*_c_ = 0.947) stretch ratios, respectively. (b) Linear correlation between stretch ratio and orientation of unmyelinated nerve fibers. Angle of 0^°^ indicates axial direction while 90^°^ indicates circumferential direction within the colorectum.

Additionally, we provide the stretch ratio for each fiber verses the fiber length (to assess dependency of the calculated stretch ratios on the length of each fiber) and representative network maps in three configurations, see Appendix A.

## 4. Discussion

In this study, we established robust and efficient methods to mechanically test, image, and analyze the deformation of networks of nerve fibers in the myenteric plexus of the colorectal tissue. By enabling precise tracking and analyses of in-plane network deformations, our methods facilitate detailed studies of fiber kinematics and mechanical properties, ultimately advancing our understanding of complex fiber networks. Our analysis methods can be applied to any two-dimensional fiber networks that can be traced in images for in-plane deformation analysis [15, 24]. This versatility allows researchers to extend the methods to a wide range of materials and structures, providing a robust tool for investigating the mechanical deformation of fibrous networks under various loading conditions [25, 26]. To facilitate application of our methods we provide free, public access to our analysis code for MATLAB, including input files for a simple test case, at github.uconn.edu/imLab/Fiber-Network_Analyses.

### 4.1. Imaging the colorectal specimen

To the best of our knowledge, this is the first attempt to conduct live fluorescent imaging of networks of nerve fibers in the colorectum subjected to high-intensity and large-strain deformations. We implemented the VGLUT2-Cre line to drive the expression of the fluorescent reporter tdTomato in most, if not all sensory neurons, which have their nerve endings concentrated at the myenteric plexus and submucosal layer [15]. We imaged the myenteric plexus from the mucosal side at a high spatial and temporal resolution to successfully capture the deformation of the complete network of nerve fibers. Crucially, our innovative design using two cylindric rods to deliver force or displacement in the circumferential direction significantly reduced through-thickness (*z*-axis) movement of the plane of the myenteric plexus, enabling in-focus imaging at a high spatial resolution during deformation.

We also kept the colorectum tubular to enable the delivery of relatively high-intensity stretch force or displacement without causing tissue failure at the fixturing due to stress concentrations. Conventional uniaxial or biaxial extension tests generally suffer from stress concentrations at the boundaries with fixturing which limits the application of force or displacement. Alternatively, inflation of the tubular colon avoids allows application of circular deformations without associated stress concentrations. However, large through-thickness displacements of the colorectum in this loading configuration pose significant challenges to clear (in-focus) imaging of the microns-thick nerve fibers, which requires a stable focal plane. We aimed to deliver high-intensity circumferential colorectal force or displacement while maintaining the planar myenteric plexus for fluorescent high-resolution imaging.

### 4.2. Analyzing the network of nerve fibers

We developed our algorithm to enable semi-automated analyses of the fiber network during tissue deformation, see Fig. 2. The heatmap of fiber stretches generated by our algorithm enables us to intuitively visualize the distribution of microns-scale fiber deformations within the network, information that could be associated with the macroscopic, tissue-level deformation. In our analyses, we defined the major range for the piece-wise fiber stretch ratio (i.e. 0.9-1.2) and separately labeled higher and lower range stretch ratios, see Fig. 3(b). We also separated fibers in tension and compression for separate analyses, allowing for more understanding of the mechanical responses and deeper insights into complex interactions within the fiber network under different loading conditions. We calculated additional data including the fiber angle with respect to the axial orientation and the fiber length. Such data enabled us to determine the potential factors contributing to variations in the stretch ratios that likely lead to variations in afferent mechanotransduction and nociception.

Our preliminary analysis of colorectal deformations indicates that the bulk-tissue deformation is larger than the deformation of the nerve fibers, a finding in agreement with current literature [27, 28]. In our representative example the tissue stretch ratio is 1.182 in the circumferential direction while the average total stretch ratio for nerve fibers is 1.048. Additionally, even the average tensile stretch ratio *λ*_t_ is 1.080, still lower than that of the bulk tissue. Our analysis of the stretch ratio with respect to the fiber angle indicates a positive slope for a linear fit, consistent with the loading conditions, see Fig 4.

### 4.3. Limitations and outlook

Naturally, there are limitations to the methods presented here. We currently determine the point cloud for analyses manually, which is relatively time consuming and doesn’t eliminate inter-user variability. In the future we aim to automate identification of the interconnection points within the fiber network, which will enhance efficiency and improve repeatability. We could achieve such automation with image processing algorithms and machine learning techniques to further refine our methods by facilitating rapid and accurate detection of network intersections [29].

In the current methods we utilized two-dimensional image stacks of VGLUT2-labeled unmyelinated nerve fibers in the myenteric plexus, an intrinsically planar anatomical structure [15, 24]. In the future we could extend our methods to accommodate three-dimensional images, which offer more comprehensive information on the morphology of fiber networks [30, 31]. Such an enhancement would significantly improve our ability to analyze and interpret the complex spatial organization and structural characteristics of nerve fibers, thereby improving our studies of mechanotransduction.

We imaged the nerve fibers using a 4*×* objective lens with a relatively large depth of field to accommodate the anticipated through-thickness (*z*-axis) movement during tissue deformation. Objective lenses with higher magnification will enhance the spatial resolution and improve detection of individual nerve fibers, but will also lead to a significant reduction in the depth of field that will cause problems with focus. To achieve higher resolution fluorescent imaging, we could implement auto-focusing algorithms to capture nerve fibers during large-strain deformations, e.g. [32].

Future studies could integrate our methods with Digital Image Correlation (DIC) to connect the deformation of the network of nerve fibers with bulk-tissue deformation under different loading modes, thus enabling comparative analyses between mechanics at macroscopic and microscopic length scales [33]. Such extended methods would facilitate comprehensive evaluation of how large-scale tissue deformations correlate with localized fiber deformations. By integrating these two levels of analysis, we can gain deeper insights into the mechanical behavior and structural responses of the fiber network, thus enhancing the accuracy and depth of our mechanistic understanding in both healthy controls and disease models.

Our methods facilitate not only improved understanding but also a structure to create, calibrate, and validated computational models, e.g. [34, 9]. Experimental data from analyses of networks of nerve fibers can provide insight on the mechanical coupling between the nerve fibers and a passive, tissue matrix, facilitating more detailed and realistic simulations of deformation behavior under mechanical loading [35]. Experimental results combined with computational models may ultimately contribute to a more comprehensive understanding of structural and functional dynamics of the nerve fibers within the colorectal environment and the subsequent visceral nociception.

## Acknowledgment

This material stems from research supported by NSF 1727185 and NIH 1R01DK120824-01.

During the preparation of this work the authors used ChatGPT-4 to sparingly edit an initial draft for brevity. After using this tool/service, the authors reviewed and edited the content in detail and take full responsibility for the content of the publication.

## Appendix A. Additional analyses

In Fig. A.5 we show supporting analyses on representative deformation of networks of nerve fibers. In Fig. A.5(a) we show the stretch ratio verses length of each fiber and conclude that there is no dependency between our calculations of stretch ratios and length of the nerve fibers. In fig. A.5(b) we show a representative fiber network at three time steps of the experiment.

**Figure A.5.**
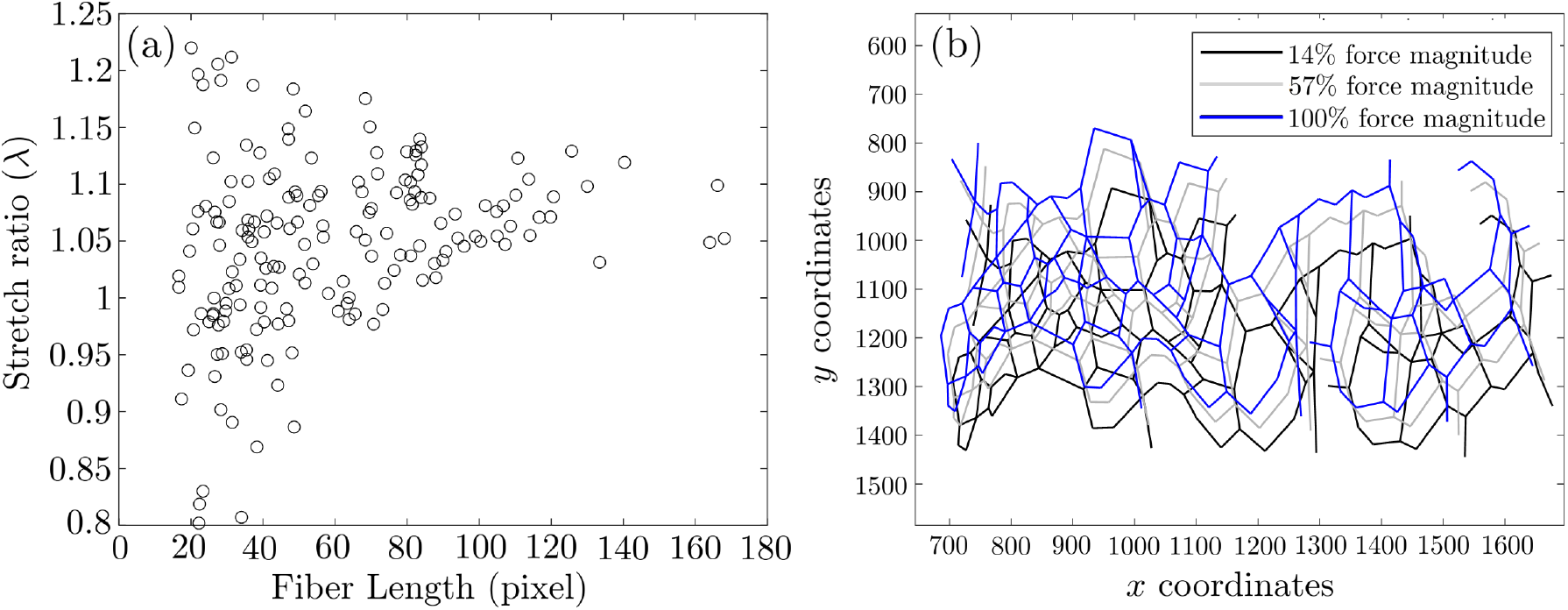
Representative supporting analyses on the deformation of networks of nerve fibers during circumferential colorectal stretch. (a) Stretch ratio verses length of each fiber. (b) Networks of nerve fibers at three time steps.

## References

[1] B. Feng, J. H. La, E. S. Schwartz, G. F. Gebhart, Irritable bowel syndrome: methods, mechanisms, and pathophysiology. neural and neuro-immune mechanisms of visceral hypersensitivity in irritable bowel syndrome, American Journal of Physiology-Gastrointestinal and Liver Physiology 302 (10) (2012) G1085–G1098.

[2] F. Cervero, J. M. Laird, Visceral pain, The Lancet 353 (9170) (1999) 2145–2148.

[3] A. Campos-Ríos, L. Rueda-Ruzafa, S. Herrera-Pérez, P. Rivas-Ramírez, J. A. Lamas, Tetrodotoxin: a new strategy to treat visceral pain?, Toxins 13 (7) (2021) 496.

[4] P. J. Pasricha, W. D. Willis, G. F. Gebhart, Chronic abdominal and visceral pain: theory and practice, CRC Press, 2006.

[5] B. Feng, P. R. Brumovsky, G. F. Gebhart, Differential roles of stretch-sensitive pelvic nerve afferents innervating mouse distal colon and rectum, American Journal of Physiology-Gastrointestinal and Liver Physiology 298 (3) (2010) G402–G409.

[6] B. Feng, T. Guo, Visceral pain from colon and rectum: the mechanotransduction and biomechanics, Journal of Neural Transmission 127 (4) (2020) 415–429.

[7] D. D. Price, Q. Zhou, B. Moshiree, M. E. Robinson, G. N. Verne, Peripheral and central contributions to hyperalgesia in irritable bowel syndrome, The Journal of Pain 7 (8) (2006) 529–535.

[8] Q. Zhou, G. N. Verne, New insights into visceral hypersensitivity—clinical implications in ibs, Nature Reviews Gastroenterology & Hepatology 8 (6) (2011) 349–355.

[9] A. Shokrani, A. Almasi, B. Feng, D. M. Pierce, Understanding mechanotransduction in the distal colon and rectum via multiscale and multimodal computational modeling, Available at SSRN 4624852.

[10] H. Eilers, M. A. Schumacher, Mechanosensitivity of primary afferent nociceptors in the pain pathway, in: A. Kamkin, I. Kiseleva (Eds.), Mechanosensitivity in Cells and Tissues, Academia, 2005.

[11] B. Feng, J.-H. La, T. Tanaka, E. S. Schwartz, T. P. McMurray, G. F. Gebhart, Altered colorectal afferent function associated with tnbs-induced visceral hypersensitivity in mice, American Journal of Physiology-Gastrointestinal and Liver Physiology 303 (7) (2012) G817–G824.

[12] N. J. Spencer, M. Kyloh, M. Duffield, Identification of different types of spinal afferent nerve endings that encode noxious and innocuous stimuli in the large intestine using a novel anterograde tracing technique, PloS one 9 (11) (2014) e112466.

[13] S. Brookes, N. Chen, A. Humenick, N. J. Spencer, M. Costa, Extrinsic sensory innervation of the gut: structure and function, The Enteric Nervous System: 30 Years Later (2016) 63–69.

[14] S. Siri, F. Maier, S. Santos, D. M. Pierce, B. Feng, Load-bearing function of the colorectal submucosa and its relevance to visceral nociception elicited by mechanical stretch, American Journal of Physiology-Gastrointestinal and Liver Physiology 317 (3) (2019) G349–G358.

[15] T. Guo, S. Patel, D. Shah, L. Chi, S. Emadi, D. M. Pierce, M. Han, P. R. Brumovsky, B. Feng, Optical clearing reveals tnbs-induced morphological changes of vglut2-positive nerve fibers in mouse colorectum, American Journal of Physiology-Gastrointestinal and Liver Physiology 320 (4) (2021) G644–G657.

[16] B. Feng, L. Chen, S. J. Ilham, A review on ultrasonic neuromodulation of the peripheral nervous system: enhanced or suppressed activities?, Applied Sciences 9 (8) (2019) 1637.

[17] J. Liu, S. Zhang, S. Emadi, T. Guo, L. Chen, B. Feng, Morphological, molecular, and functional characterization of mouse glutamatergic myenteric neurons, American Journal of Physiology-Gastrointestinal and Liver Physiology 326 (3) (2024) G279–G290.

[18] T. Guo, Z. Bian, K. Trocki, L. Chen, G. Zheng, B. Feng, Optical recording reveals topological distribution of functionally classified colorectal afferent neurons in intact lumbosacral drg, Physiological reports 7 (9) (2019) e14097.

[19] T. Guo, J. Liu, L. Chen, Z. Bian, G. Zheng, B. Feng, Sex differences in zymosan-induced behavioral visceral hypersensitivity and colorectal afferent sensitization, American Journal of Physiology-Gastrointestinal and Liver Physiology 326 (2) (2024) G133–G146.

[20] J. Humphrey, S. DeLange, An Introduction to Biomechanics: Solids and Fluids, Analysis and Design, Springer Link, 2015.

[21] A. D. Edelstein, M. A. Tsuchida, N. Amodaj, H. Pinkard, R. D. Vale, N. Stuurman, Advanced methods of microscope control using μmanager software, Journal of biological methods 1 (2).

[22] J. Schindelin, I. Arganda-Carreras, E. Frise, V. Kaynig, M. Longair, T. Pietzsch, S. Preibisch, C. Rueden, S. Saalfeld, B. Schmid, et al., Fiji: an open-source platform for biological-image analysis, Nature methods 9 (7) (2012) 676–682.

[23] P.-E. Danielsson, Euclidean distance mapping, Computer Graphics and image processing 14 (3) (1980) 227–248.

[24] T. Guo, S. Patel, D. Shah, L. Chi, S. Emadi, D. M. Pierce, M. Han, P. R. Brumovsky, B. Feng, Neurogastroenterology and motility: Optical clearing reveals tnbs-induced morphological changes of vglut2-positive nerve fibers in mouse colorectum, American Journal of Physiology-Gastrointestinal and Liver Physiology 320 (4) (2021) G644.

[25] E. Sozumert, V. Cucumazzo, V. V. Silberschmidt, 9 - deformation and damage of random fibrous networks, in: V. V. Silberschmidt (Ed.), Mechanics of Fibrous Networks, Elsevier Series in Mechanics of Advanced Materials, Elsevier, 2022, pp. 203–219.

[26] E. Sozumert, V. V. Silberschmidt, 1 - mechanics of fibrous networks: Basic behaviour, in: V. V. Silberschmidt (Ed.), Mechanics of Fibrous Networks, Elsevier Series in Mechanics of Advanced Materials, Elsevier, 2022, pp. 1–12.

[27] A. Tamura, K. Nagayama, T. Matsumoto, Measurement of nerve fiber strain in brain tissue subjected to uniaxial stretch (comparison between local strain of nerve fiber and global strain of brain tissue), Journal of Biomechanical Science and Engineering 1 (2) (2006) 304–315.

[28] A. Tamura, K. Nagayama, T. Matsumoto, S. Hayashi, Variation in nerve fiber strain in brain tissue subjected to uniaxial stretch, Tech. rep., SAE Technical Paper (2007).

[29] J. R. Parker, Algorithms for image processing and computer vision, John Wiley & Sons, 2010.

[30] Y.-A. Liu, Y.-C. Chung, S.-T. Pan, Y.-C. Hou, S.-J. Peng, P. J. Pasricha, S.-C. Tang, 3-d illustration of network orientations of interstitial cells of cajal subgroups in human colon as revealed by deep-tissue imaging with optical clearing, American Journal of Physiology-Gastrointestinal and Liver Physiology 302 (10) (2012) G1099–G1110.

[31] Y. Liu, Y. Chung, S. Pan, M. Shen, Y. Hou, S. Peng, P. Pasricha, S. Tang, 3-d imaging, illustration, and quantitation of enteric glial network in transparent human colon mucosa, Neuro-gastroenterology & Motility 25 (5) (2013) e324–e338.

[32] Y. Sun, S. Duthaler, B. J. Nelson, Autofocusing algorithm selection in computer microscopy, in: 2005 IEEE/RSJ International Conference on Intelligent Robots and Systems, IEEE, 2005, pp. 70–76.

[33] B. Pan, Recent progress in digital image correlation, Experimental mechanics 51 (2011) 1223–1235.

[34] Y. Zhao, S. Siri, B. Feng, D. M. Pierce, The macro-and micro-mechanics of the colon and rectum ii: theoretical and computational methods, Bioengineering 7 (4) (2020) 152.

[35] S. Kakaletsis, E. Lejeune, M. Rausch, The mechanics of embedded fiber networks, Journal of the Mechanics and Physics of Solids 181 (2023) 105456.

